# A framework for testing different imputation methods for tabular datasets

**DOI:** 10.1101/773762

**Authors:** Tabea Kossen, Michelle Livne, Vince I Madai, Ivana Galinovic, Dietmar Frey, Jochen B Fiebach

## Abstract

**Background and purpose:** Handling missing values is a prevalent challenge in the analysis of clinical data. The rise of data-driven models demands an efficient use of the available data. Methods to impute missing values are thus crucial. Here, we developed a publicly available framework to test different imputation methods and compared their impact in a typical stroke clinical dataset as a use case.

**Methods:** A clinical dataset based on the 1000Plus stroke study with 380 completed-entries patients was used. 13 common clinical parameters including numerical and categorical values were selected. Missing values in a missing-at-random (MAR) and missing-completely-at-random (MCAR) fashion from 0% to 60% were simulated and consequently imputed using the mean, hot-deck, multiple imputation by chained equations, expectation maximization method and listwise deletion. The performance was assessed by the root mean squared error, the absolute bias and the performance of a linear model for discharge mRS prediction.

**Results:** Listwise deletion was the worst performing method and started to be significantly worse than any imputation method from 2% (MAR) and 3% (MCAR) missing values on. The underlying missing value mechanism seemed to have a crucial influence on the identified best performing imputation method. Consequently no single imputation method outperformed all others. A significant performance drop of the linear model started from 11% (MAR+MCAR) and 18% (MCAR) missing values.

**Conclusions:** In the presented case study of a typical clinical stroke dataset we confirmed that listwise deletion should be avoided for dealing with missing values. Our findings indicate that the underlying missing value mechanism and other dataset characteristics strongly influence the best choice of imputation method. For future studies with similar data structure, we thus suggest to use the developed framework in this study to select the most suitable imputation method for a given dataset prior to analysis.

## Introduction

Missing values are a prevalent challenge in the analysis of clinical data [1–3]. Validated guidelines for handling missing data are more important now than ever with the rise of data-driven applications [4–8]. Here, an efficient utilization of the data is crucial in the limited settings of usually fairly small medical datasets [9]. Additionally, while deletion of patient entries with missing values (listwise deletion) is a common practice, it can lead to biased results and is therefore highly discouraged [1, 10, 11].

An alternative approach is to apply imputation methods on the missing data [12, 13]. Imputation methods allow replacing missing values with substituted values that estimate the true underlying value. Most commonly used methods in clinical datasets include simple imputation methods like mean imputation and hot-deck imputation (“sampling”) as well as more complex algorithms like multiple imputation by chained equations (MICE) [14] and expectation maximization (EM) using multiple imputation [15].

However, there is a controversy regarding which imputation method should be used [12, 16, 17]. More importantly, only few studies exist which assessed imputation methods in the medical field [13, 16–18]. This is also true in the stroke field. While Young-Saver et al. investigated the imputation of stroke outcome data, to date no study has compared and validated imputation methods for a typical clinical stroke dataset as a whole [19].

The objectives of this work were thus to 1) develop a publicly available framework (https://github.com/tabeak/missing-value-analysis) which can identify the best imputation method for a given tabular dataset and 2) to compare different imputation method for handling missing data in clinical stroke dataset as a use case for the framework. When comparing the different imputation methods, we assessed both how much the imputed values differed from the ground truth as well as how the different imputation methods influenced the performance of a data-driven predictive model.

## Materials and methods

### Patients

A clinical dataset based on the 1000Plus stroke study with 380 completed-entries of acute stroke patients was used [20]. The study was approved by the institutional review board of the Charité Universitätsmedizin Berlin. All patients gave their written informed consent. Because of the sensitive nature of the data collected for this study, requests to access the dataset from qualified researchers trained in human subject confidentiality protocols may be sent to the institutional ethics commitee of Charité Universitätsmedizin Berlin.

### Data analysis

13 common clinical parameters were analyzed. Among them, seven were numerical: hours-to-MRI, age, pre-stroke mRS, acute National Institutes of Health Stroke Scale (NIHSS), discharge NIHSS, discharge mRS and discharge Trial-of-ORG-10172-in-Acute-Stroke-Treatment (TOAST). Six were categorical: sex, treatment with tissue plasminogen activator (tPA), occlusion, hyperlipidemia, diabetes and hypertonia.

Missing values from 0% to 60% were simulated following two different cases of missing values: missing-at-random (MAR) and missing-completely-at-random (MCAR). MAR means that the probability for a value missing depends on same values of other observed variables. MCAR, on the other hand, describes the scenario that values are missing completely at random. In contrast to MAR, there is no systematic reason for a missing value and the probability for a value missing is the same for each value. Performance was estimated for different imputation methods in two fashions: 1) Error assessment using RMSE and absolute bias and 2) Performance assessment of stroke discharge mRS. The imputation methods included 1) mean imputation, 2) hot-deck imputation, 3) MICE and 4) multiple imputation by EM and 5) listwise deletion.

#### Error assessment

For the error assessment analysis, we chose two common measures for evaluating imputation methods, RMSE and absolute bias [21]. In this analysis, the parameters were split into numerical and categorical. For numerical parameters, the RMSE of the normalized data is defined according to:

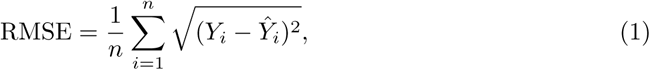

where *n* is the number of imputed samples, *Ŷ*_*i*_ the estimated sample value and *Y*_*i*_ the true value. For categorical parameters, the RMSE corresponds to the percentage of misclassified values:

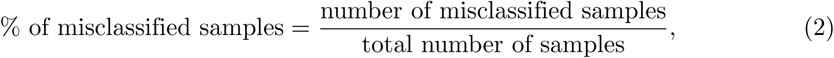

As a second error assessment, the mean absolute bias was calculated. It is defined as:

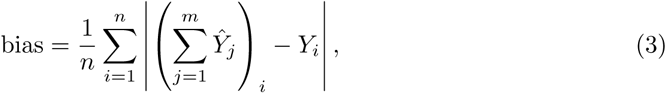

where *m* is the number of iterations for each imputation, i.e. how often the value was imputed. The absolute bias was then averaged over all imputed samples *n*.

Both RMSE as well as the absolute bias was assessed for each variable at a time and then averaged for each parameter-type (i.e. numerical vs. categorical).

#### Predictive model analysis

The second part of the analysis included the incorporation of a supervised predictive model, the generalized linear model (GLM). We constructed a model predicting the modified Rankin Scale (mRS) at discharge, which is a measure of the early clinical outcome after stroke. It can take values from 0 (no symptoms) to 6 (death). The mRS was split into 0-2 (good outcome) and 3-6 (bad outcome) [4, 22]. Other discharge parameters (discharge NIHSS and discharge TOAST) were excluded for this analysis to maintain the integrity of the model. This predictive modelling framework is used as a standard method for this use case [23–25]. As the MAR mechanism deletes parameter values with respect to other parameter’s values, the mechanism could only be simulated for up to 9% missing values. Therefore, missing values from 10% to 60% are deleted combining the MAR mechanism with the MCAR mechanism. For simplicity, this mechanism is hence termed “MAR+MCAR” throughout this work.

Receiver operating characteristics (ROC) analysis was applied to assess the model performance and the imputation methods were then compared based on the area under the curve (AUC). Further, the methodologies were compared to listwise deletion. Listwise deletion removes every patient that has at least one missing value and thus, reduces the size of the dataset. Due to this limitation, the GLM could only be computed for up to 10% missing values for this sub-analysis.

### Statistical analysis

Both the error and the predictive model analyses were repeated 100 times for randomly simulated missing values. For the RMSE, each imputation method was compared to each of the other methods using the paired Wilcoxon signed-rank test for each respective percentage value. The same analysis was conducted for performance assessment of the predictive model. To calculate the mean absolute bias, the 100 repeated imputations are averaged and then compared to the true value. This results in one value per percentage. Thus, a statistical analysis cannot be performed for each percentage.

Finally, the threshold of the percentage of missing values to significantly impair performance was determined for the different imputation methods: The performance of the completed-entry dataset (0% missing values) was compared to the performance for each percentage of missing value using the paired Wilcoxon signed-rank test. The threshold was identified when the performance drop was found to be significant according to the standard value of *p* < 0.05.

## Results

### Clinical data

380 acute stroke patients had complete entries for the 13 clinical parameters. The median age was 72 and the median NIHSS score was 3. The distribution of the clinical parameters is given in Table 1.

**Table 1.** Original distribution of clinical parameters.

| Clinical parameter | Value |
| --- | --- |
| median hours-to-MRI (IQR) | 11 (15) |
| median age (IQR) | 72 (15) |
| median mRS prestroke (IQR) | 0 (1) |
| median acute NIHSS (IQR) | 3 (4.25) |
| median discharge NIHSS (IQR) | 1 (3) |
| median discharge mRS (IQR) | 1 (2) |
| median discharge TOAST (IQR) | 1 (1) |
| females (%) / males (%) | 247 (65) / 133 (35) |
| tPA treatment yes / no (%) | 93 (24.47) / 287 (75.53) |
| occlusion yes / no (%) | 219 (57.63) / 161 (42.37) |
| hyperlipidemia yes / no (%) | 224 (58.95) / 156 (41.05) |
| diabetes yes / no (%) | 286 (75.26) / 94 (24.74) |
| hypertonia yes / no (%) | 309 (81.32) / 71 (18.68) |

Median values and IQR of the numerical clinical parameters hours-to-MRI, age, mRS pre-stroke, acute NIHSS, discharge NIHSS, discharge mRS and discharge TOAST. Total amount of patients and percentages of the categorical parameters sex, treatment with tPA, occlusion, hyperlipidemia, diabetes and hypertonia. (Abbreviations: IQR = interquartile range, MRI = magnetic resonance imaging, mRS = modified Rankin scale, NIHSS = National Institutes of Health Stroke Scale, TOAST = Trial-of-ORG-10172-in-Acute-Stroke-Treatment, tPA = tissue plasminogen activator).

### Error assessment

Figs 1 to 4 show the RMSE and absolute bias for increasing MAR and MCAR missing values. The figures are separated into the averaged and the accumulated values of both numerical (Figs 1 and 4) and categorical data (Figs 2 and 4). While the accumulated error increases with larger percentages of missing values, the averaged RMSE and absolute bias remains relatively steady. No single imputation method consistently outperformed all other methods.

**Fig 1.**
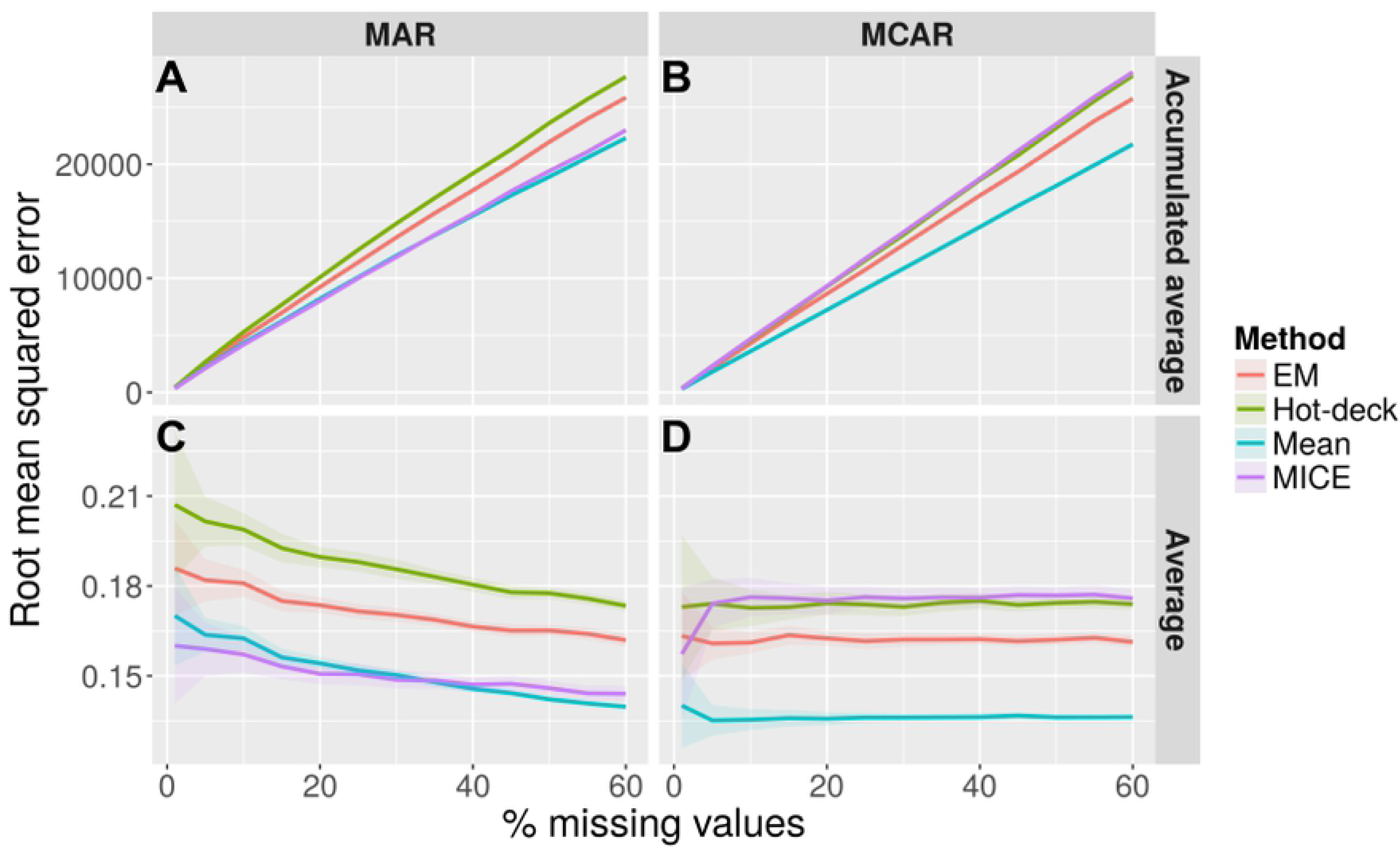
Mean RMSE for increasing percentages of missing data for numerical parameters. Mean RMSE for increasing missing values from 0% to 60% using mean imputation (blue), hot-deck imputation (olive), MICE (purple) and EM (red) for numerical data. (A) describes the accumulated average RMSE if missing values are generated in a MAR fashion, (B) in a MCAR fashion. (C) shows the average RMSE if missing values are generated using the MAR mechanism and (D) the MCAR mechanism. (Abbreviations: EM = expectation maximization, MAR = missing-at-random, MCAR = missing-completely-at-random, MICE = multiple imputation by chained equations, RMSE = root mean squared error)

**Fig 2.**
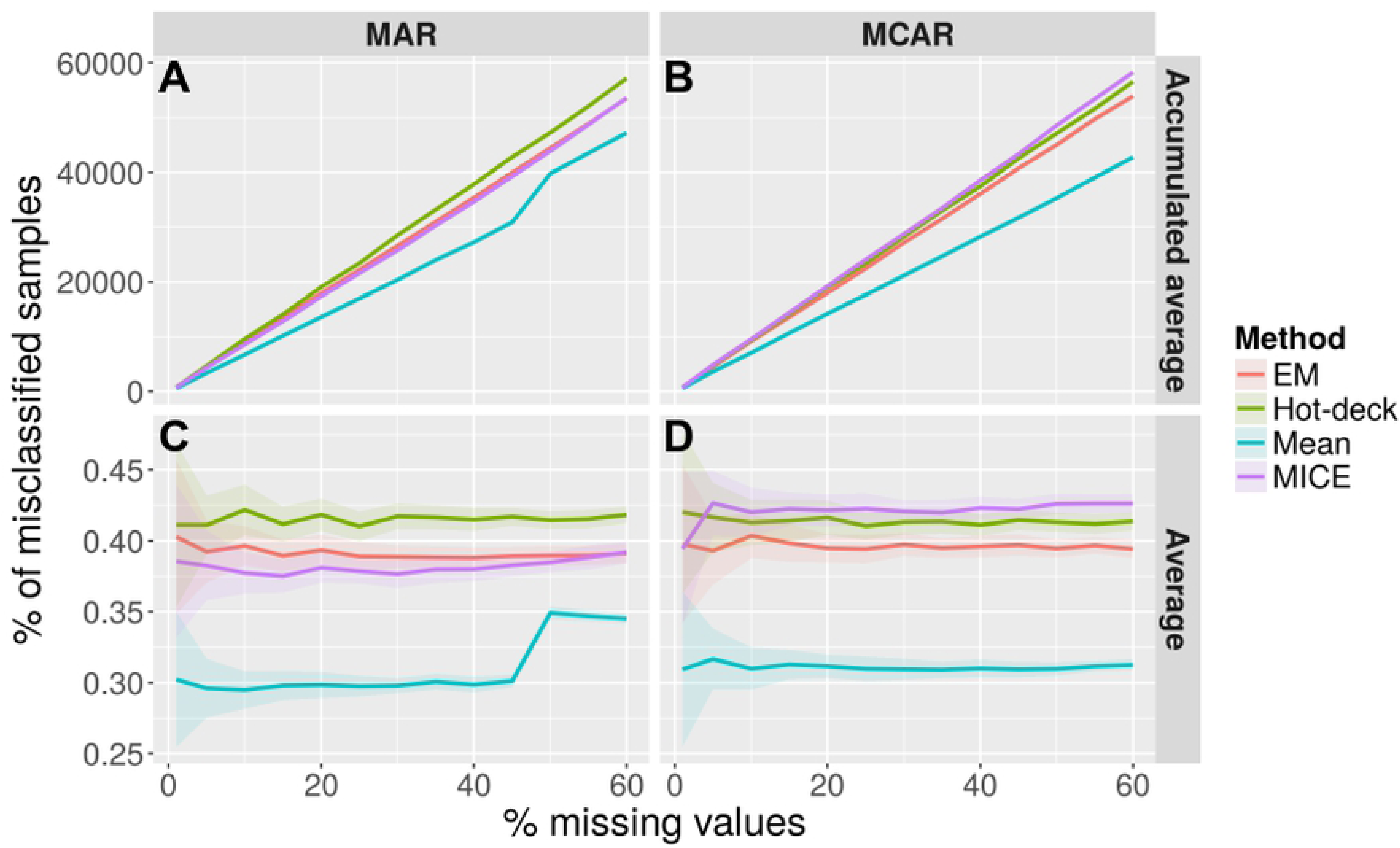
Mean percentage of misclassified samples for increasing percentages of missing data for categorical parameters. Mean percentage of misclassified samples for increasing missing values from 0% to 60% using mean imputation (blue), hot-deck imputation (olive), MICE (purple) and EM (red) for categorical data. (A) describes the accumulated average percentage of misclassified samples if missing values are generated in a MAR fashion, (B) in a MCAR fashion. (C) shows the average percentage of misclassified samples if missing values are generated using the MAR mechanism and (D) the MCAR mechanism. (Abbreviations: EM = expectation maximization, MAR = missing-at-random, MCAR = missing-completely-at-random, MICE = multiple imputation by chained equations)

**Fig 3.**
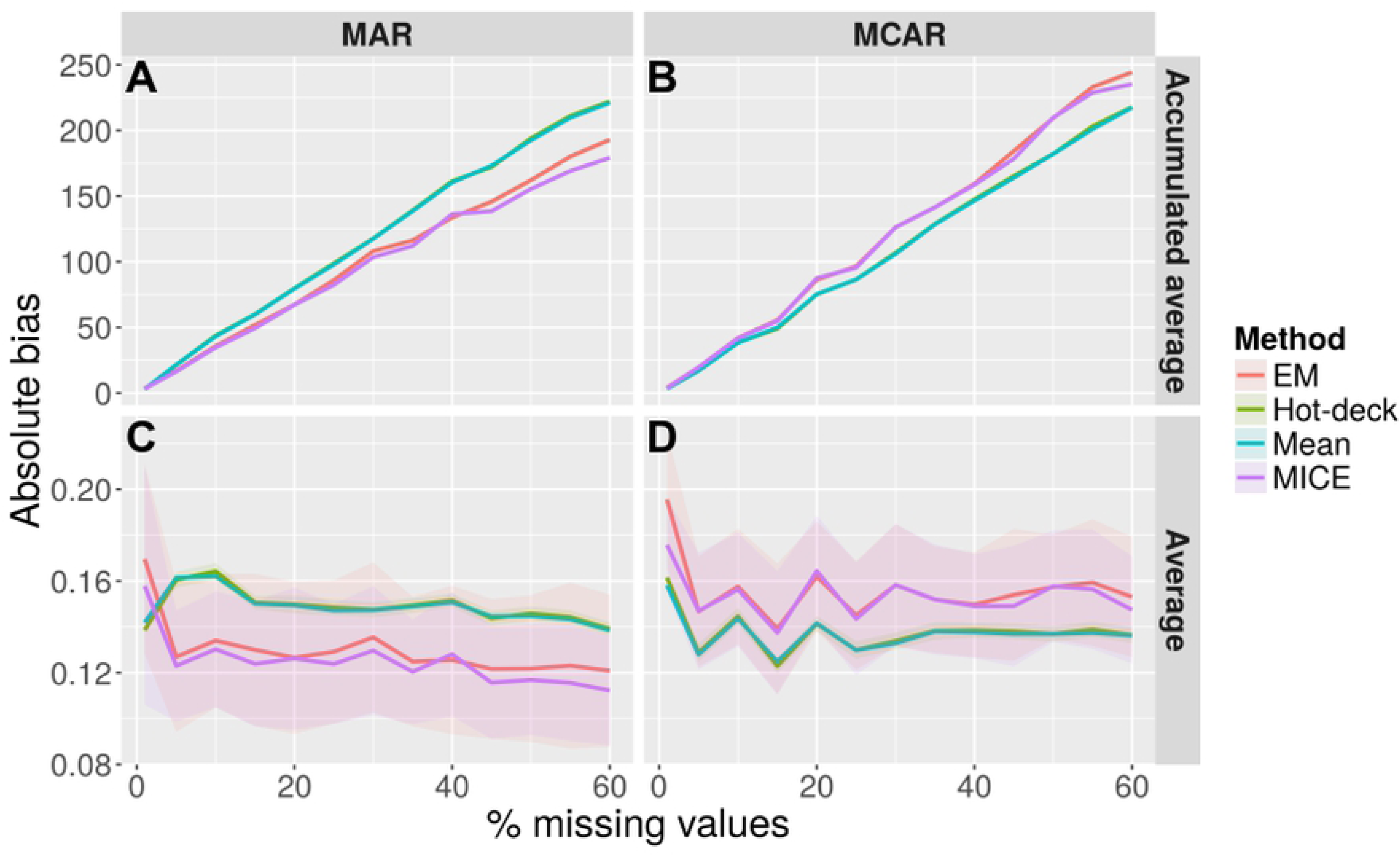
Mean absolute bias for increasing percentages of missing data for numerical parameters. Mean absolute bias for increasing missing values from 0% to 60% using mean imputation (blue), hot-deck imputation (olive), MICE (purple) and EM (red) for numerical data. (A) describes the accumulated average bias if missing values are generated in a MAR fashion, (B) in a MCAR fashion. (C) shows the average bias if missing values are generated using the MAR mechanism and (D) the MCAR mechanism. Note that the error margin in (C) and (D) corresponds to the standard deviation of the samples estimates and not the bias. (Abbreviations: EM = expectation maximization, MAR = missing-at-random, MCAR = missing-completely-at-random, MICE = multiple imputation by chained equations)

**Fig 4.**
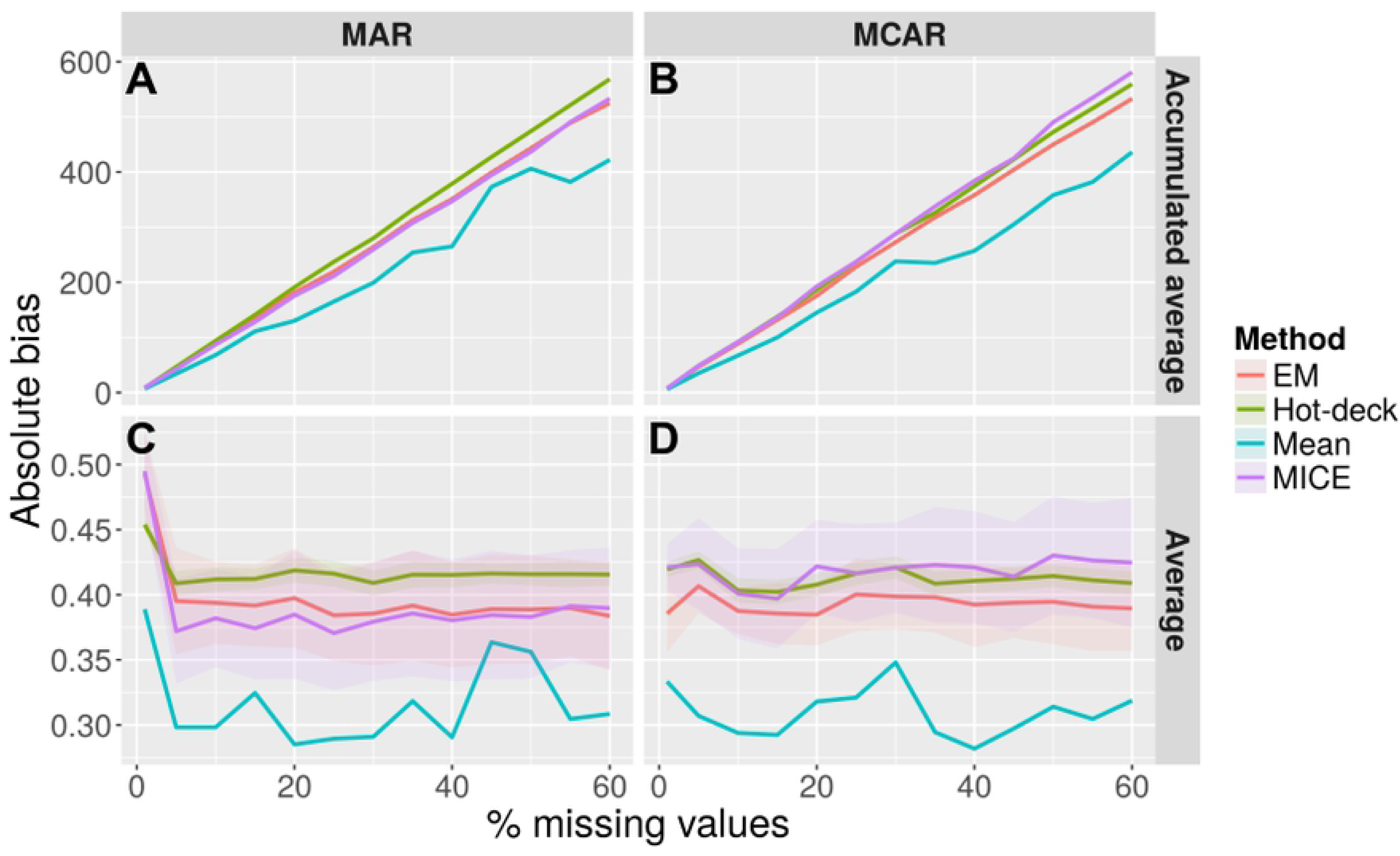
Mean absolute bias for increasing percentages of missing data for categorical parameters. Mean absolute bias for increasing missing values from 0% to 60% using mean imputation (blue), hot-deck imputation (olive), MICE (purple) and EM (red) for categorical data. (A) describes the accumulated average bias if missing values are generated in a MAR fashion, (B) in a MCAR fashion. (C) shows the average bias if missing values are generated using the MAR mechanism and (D) the MCAR mechanism. Note that the error margin in (C) and (D) corresponds to the standard deviation of the samples estimates and not the bias. (Abbreviations: EM = expectation maximization, MAR = missing-at-random, MCAR = missing-completely-at-random, MICE = multiple imputation by chained equations)

The RMSE is shown in Figs 1 and 2. For the numerical data MICE and mean imputation show the lowest error rate for MAR missing values and mean imputation for MCAR missing values (Fig 1). For the categorical data the lowest percentage of misclassified samples could be observed for mean imputation (Fig 2). For MAR missing values the mean imputation appears less steady and stable compared to the MCAR missing values.

Both mean imputation and MICE show a significantly lower RMSE than hot-deck and EM for numerical data in the MAR type-case (*p* < 0.05). For MCAR missing values as well as MAR missing values on categorical data mean imputation in terms of RMSE performed significantly better than the other imputation methods (*p* < 0.05).

Figs 3 and 4 show the absolute bias. For numerical data MICE and EM showed the lowest absolute bias in the MAR case and mean and hot-deck imputation in the MCAR case. Mean imputation showed the lowest bias for categorical data for both MAR and MCAR. The mean imputation yielded less stable results for categorical data compared to numerical data.

### Predictive model analysis

Fig 5 shows the performance of the GLM for an increasing amount of missing values that were generated in a MAR+MCAR fashion. Generally, the more values were imputed, the lower the resulting performance. The best overall performance was yielded by mean imputation (Fig 5A). Compared to the other imputation methods, this difference is significant only in the range of 25% to 45% missing values (Fig 5C). Listwise deletion showed the lowest performance compared to all other imputation methods (Fig 5B). Starting from 2% missing values every other imputation method performed significantly better than listwise deletion (Fig 5D).

**Fig 5.**
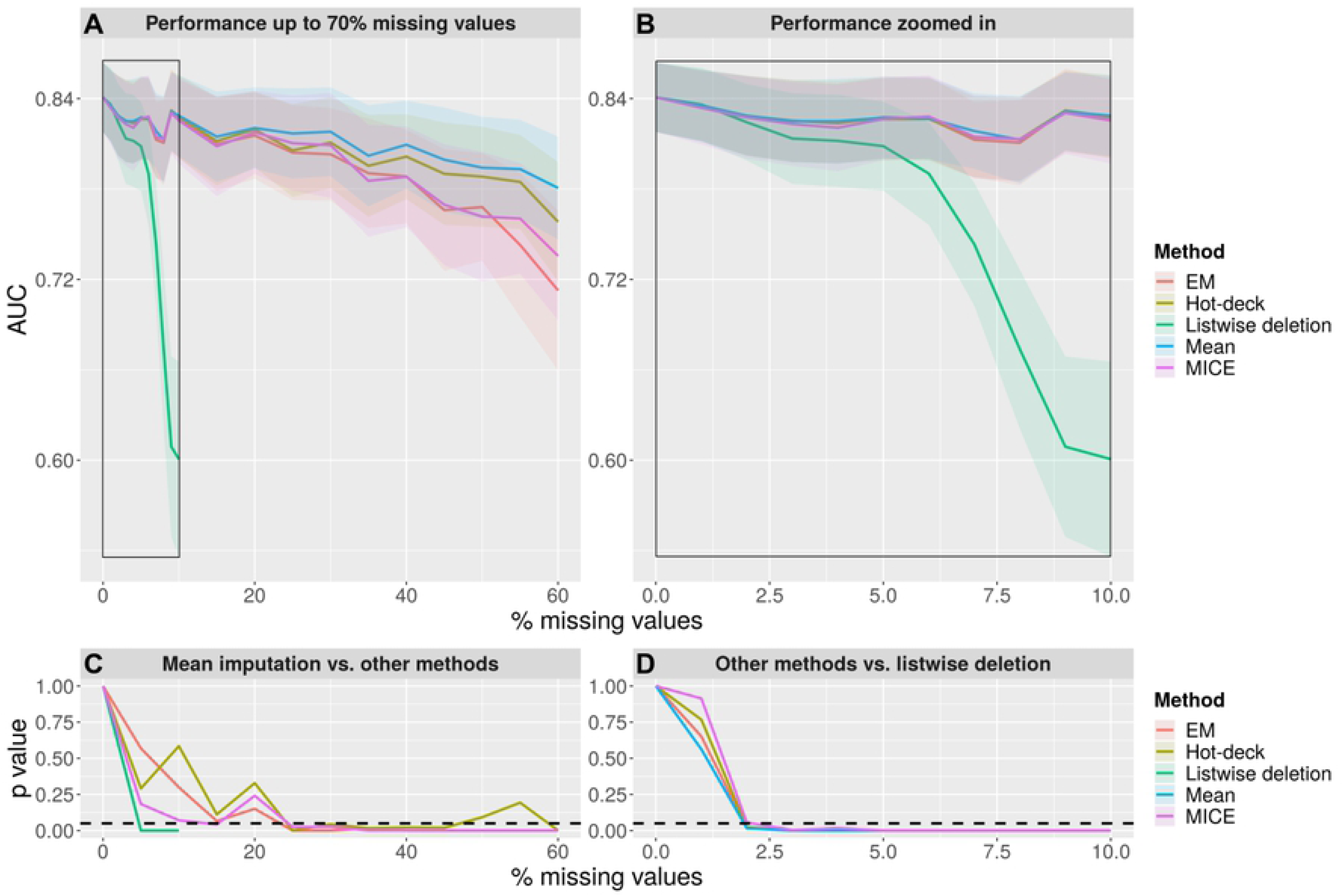
GLM performance on the dataset with increasing percentages of MAR+MCAR missing values and comparison of imputation methods with listwise deletion and mean imputation. (A) and (B) show the predictive model performance in terms of AUC for increasing MAR+MCAR missing values from range 0% to 60% and 0% to 10% respectively using mean (blue), hot-deck imputation (olive), MICE (purple), EM imputation (red) and listwise deletion (green). The plots in the bottom (C) and (D) show the corresponding *p* values of the different imputation methods compare to mean imputation (C) and listwise deletion (D) using a paired Wilcoxon signed-rank test. The horizontal dashed black line indicates 0.05, the threshold of significance for the *p* values. (Abbreviations: AUC = area under the curve, GLM = generalized linear model, EM = expectation maximization, MAR = missing-at-random, MICE = multiple imputation by chained equations)

The completed-entry model (0% missing values) showed higher AUC values compared to all imputation methods. The difference started to be significant for the MAR case-type between 2% to 3% missing values. From 11% on every model is significantly worse than the complete-entry model.

Similar results could be observed for MCAR missing values (Fig 6). The more values imputed, the lower the resulted AUC is. Mean imputation yielded the best performance, yet significance was shown only for 45% missing values and above (Figs 6A and 6C). Listwise deletion performed significantly lower than all other imputation methods starting from 3% missing values (Fig 6D).

**Fig 6.**
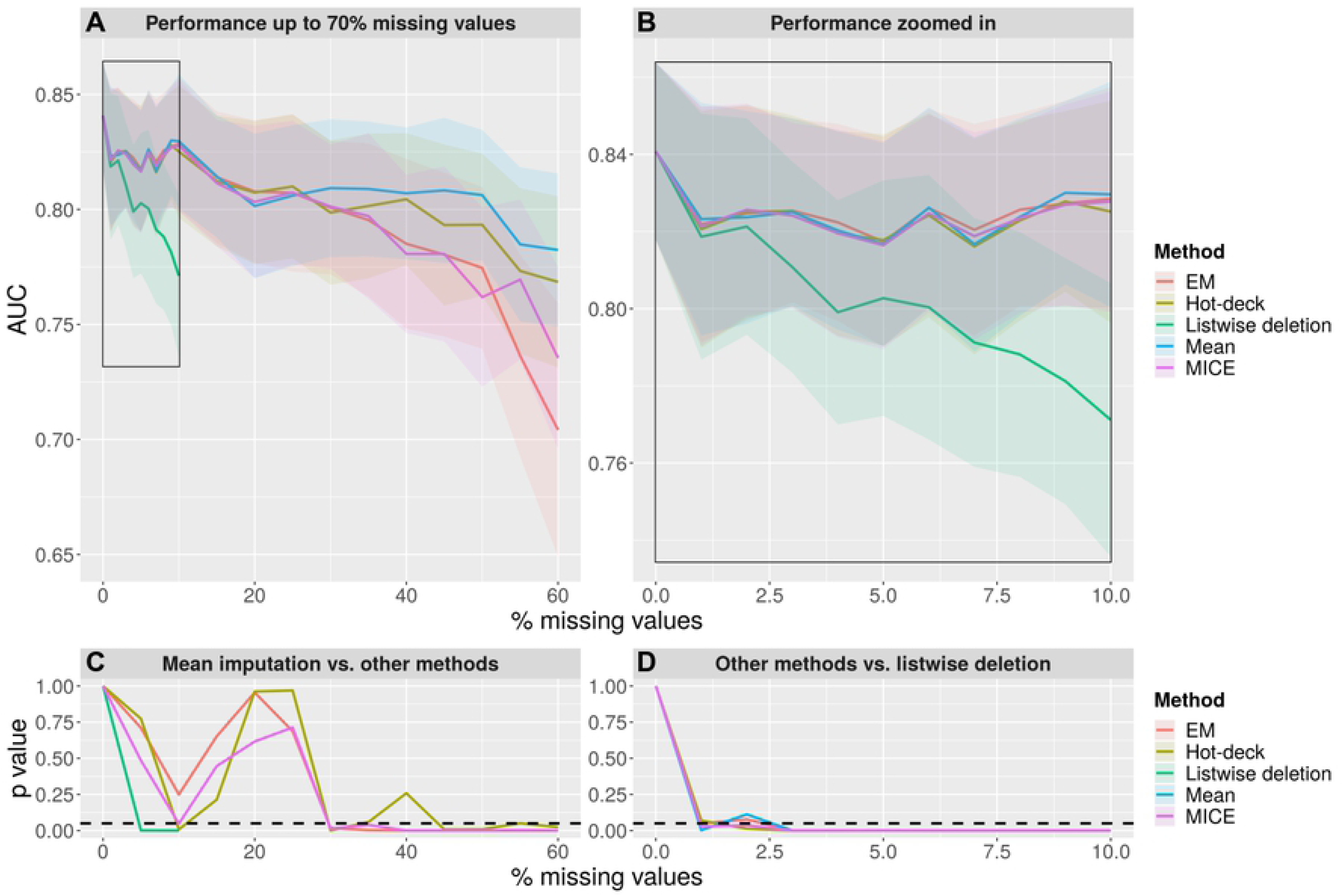
GLM performance on the dataset with increasing percentages of MCAR missing values and comparison of imputation methods with listwise deletion and mean imputation. (A) and (B) show the predictive model performance in terms of AUC for increasing MCAR missing values from range 0% to 60% and 0% to 10% respectively using mean (blue), hot-deck imputation (olive), MICE (purple), EM imputation (red) and listwise deletion (green). The plots in the bottom (C) and (D) show the corresponding *p* values of the different imputation methods compare to mean imputation (C) and listwise deletion (D) using a paired Wilcoxon signed-rank test. The horizontal dashed black line indicates 0.05, the threshold of significance for the *p* values. (Abbreviations: AUC = area under the curve, GLM = generalized linear model, EM = expectation maximization, MCAR = missing-completely-at-random, MICE = multiple imputation by chained equations)

For the MCAR case-type, the completed-entry model performed the best as well. The first significant AUC value was for 1% missing values. Starting from 18% every model was significantly worse than the complete-entry model.

## Discussion

In the present study, we developed a publicly available framework to investigate different imputation methods for handling missing values and tested it in a clinical stroke dataset as a use case. The utilized dataset 1000Plus represents a typical dataset in stroke regarding size and recorded values. For the predictive model, the results show that listwise deletion performs significantly worse than imputation methods starting from a low percentage (2% for MAR and 3% for MCAR). Additionally, our results indicate that for this type of data you should not impute data above 10% (MAR+MCAR) and 17% (MCAR). For the error assessment no method outperformed all other methods for every analysis. Furthermore, it seems to be crucial which missing value mechanism is underlying in the dataset.

Listwise deletion is still commonly practiced yet highly discouraged [1, 10]. Our results corroborate this notion and strongly suggest to use imputation methods. The performance of our predictive model started to drop significantly already when only 2% of the values were missing using listwise deletion. This implies that the available incomplete patient information still adds crucial value to the predictive model and should not be neglected.

Our results do not provide a strict recommendation for one imputation method. While mean imputation seemed to show the lowest RMSE and highest performance in terms of AUC, these results should be interpreted with caution. Mean imputation is a method that aims to reduce the RMSE, thus this measurement is biased towards mean imputation. Therefore, we additionally compared the methodologies using the absolute bias. Here, mean imputation performs well for categorical data as well as numerical data with MCAR missing values. Looking at the error assessment for categorical data, however, we observed that mean imputation performed less robustly. In the particular case of categorical data, mean imputation means imputing the value that occurs most often in the remaining dataset. Hence, the imputation method highly depends on which category the missing value belonged to. The resulting error is then less stable and more easily corrupted by the missing value pattern.

In the predictive model analysis, mean imputation showed significantly better results than other imputation methods in the range of 25% to 45% (MAR+MCAR) and 45% to 60% (MCAR) missing values. For the given dataset we establish a threshold of 11% (MAR+MCAR) and 18% (MCAR) over which imputation of missing values is discouraged. Consequently, the significant improvement of mean imputation is a priori not within the practical range where values should be imputed [26, 27].

For numerical data in the MAR case-type, we found MICE and EM to show the lowest absolute bias. In other studies, complex algorithms like MICE and EM also appeared to be superior to seemingly old-fashioned imputation methods like mean or hot-deck imputation [16, 17, 26]. In the case of numerical data and MCAR missing values, however, mean and hot-deck imputation showed the lowest bias. It seems unintuitive that simple algorithms like mean and sampling as the best performing imputation methods. The higher bias for MCAR compared to simpler imputation methods might, however, be explained by inherent characteristics of the more sophisticated methods. The MICE algorithm builds upon strong dependencies between the covariates. The missing value is estimated based on the corresponding values of the other parameters. When removing values in a completely random fashion, i.e. MCAR, the dependencies between the covariates might not be as strong anymore in our dataset. Thus, it is hard to reconstruct. If values are missing in a non-completely random fashion, i.e. MAR, there is pattern for missingness available that complex algorithms like MICE can learn from. The same holds true for the EM. The EM algorithm estimates the underlying log likelihood of the complete dataset [28, 29]. Given this distribution, the missing values are approximated. In the MAR case unlike MCAR, the existing pattern of missing values might help to capture the likelihood to yield a good estimation. To conclude, the underlying missing value mechanism might be very crucial regarding which imputation method is the most suitable for the given dataset.

In the error assessment, we observed that the error rate was quite constant for an increasing percentage of missing data. While appearing counterintuitive on first sight, the explanation for this phenomenon is simple: For each imputation method, we assess a value from a distribution that estimates the true underlying distribution. When drawing an increasing number of samples from both distributions, the difference between the values drawn from the two distributions remains the same on average. Thus, the error rate does not increase for more missing samples. Nevertheless, when looking at the accumulated error, we can see that for each imputed sample the new error is added so that the error rate is in fact increasing.

In the predictive model performance assessment, however, we see a different behavior. With increasing amount of missing values we can see decreasing predictive performance [30–32]. Depending on the size of the dataset and the number of covariates, the performance drops at a certain threshold. Thus, the significant decrease in performance can occur at a different percentage of missing values. In our study with 380 patients and 10 predictive parameters, the significant performance drop was measured starting from 11% missing values for MAR+MCAR missing values and 18% for MCAR respectively. Therefore, we suggest to only use imputation methods until 10% missing values. Our results confirm findings from the literature identifying numbers in a similar range of missing value percentages leading to a performance drop [26, 27].

Importantly, our results implicate that there is no generally “best” imputation method. Our findings suggest that – under certain circumstances – simple mean imputation might be superior to the other sophisticated imputation techniques. In other cases, i.e. MAR as the underlying missing value mechanism, MICE or EM performed better. This is corroborated also in theory by the “no free lunch theorem” [33, 34]. The theorem states that there is no algorithm that performs best in all tasks. The good performance of one algorithm in one task comes with the cost of low performance in another task [33]. Since the imputation methods are in fact algorithms and the different dataset can be seen as tasks, the theorem could apply here as well. Hence, our results are specific for our dataset. Distinct characteristics of any other given dataset like its size, mechanism of data missingness and the type of features will influence which imputation method should be preferred. Thus, we make our framework publicly available (https://github.com/tabeak/missing-value-analysis). It can easily be used and adapted by other researchers to test their own datasets and identify the optimal imputation method for their data. Especially given the often limited size of datasets in medical applications, such an approach might allow increasing the validity of statistical testing and predictive modeling. Finally, our work is strongly encouraging further research on the performance of imputation methods in other tabular datasets.

Our study has several limitations. First of all, we could simulate MAR missing values only up to 9% due to mathematical constraints on covariates dependencies and the limited size of our dataset. Hence, our analysis mostly relates to mixed MAR+MCAR and MCAR mechanisms. The real underlying mechanism for missing values in clinical stroke datasets remains unknown. It is likely, however, that the true missing data mechanism is a mixture of MAR and MCAR as missing values can occur systematically as well as randomly in medical datasets. Secondly, due to data availability, we trained our predictive model on discharge mRS and not on final three months mRS, which is the clinically more useful measure. However, given the methodological nature of our study, the predictive model is only exemplary to show the impact of different methods dealing with missing data. The impact of missing data and different imputation methods on models predicting three months mRS must be elucidated in future studies.

## Conclusion

We developed a publicly available R framework to evaluate different imputation methods and tested it on a typical clinical stroke dataset as a use case. Our main finding was that listwise deletion should not be performed and the choice of imputation methods might depend highly on the underlying missing value mechanism and other characteristics of a given dataset. Thus, we suggest that the optimal imputation method is dataset-dependent and we strongly encourage other researchers to adapt our openly available framework to their own datasets prior to analysis.

